# SpliceAI-10k calculator for the prediction of pseudoexonization, intron retention, and exon deletion

**DOI:** 10.1101/2022.07.30.502132

**Authors:** Daffodil M. Canson, Aimee L. Davidson, Miguel de la Hoya, Michael T. Parsons, Dylan M. Glubb, Olga Kondrashova, Amanda B. Spurdle

## Abstract

**Summary:** SpliceAI is a widely used splicing prediction tool and its most common application relies on the maximum delta score to assign variant impact on splicing. We developed the SpliceAI-10k calculator (SAI-10k-calc) to extend use of this tool to predict: the splicing aberration type including pseudoexonization, intron retention, partial exon deletion, and (multi)exon skipping using a 10 kb analysis window; the size of inserted or deleted sequence; the effect on reading frame; and the altered amino acid sequence.

SAI-10k-calc has 95% sensitivity and 96% specificity for predicting variants that impact splicing, computed from a control dataset of 1,212 single nucleotide variants (SNVs) with curated splicing assay results. Notably, it has high performance (≥84% accuracy) for predicting pseudoexon and partial intron retention. The automated amino acid sequence prediction allows for efficient identification of variants that are expected to result in mRNA nonsense-mediated decay or translation of truncated proteins.

**Availability and implementation:** SAI-10k-calc is implemented in R (https://github.com/adavi4/SAI-10k-calc) and also available as a Microsoft Excel spreadsheet. Users can adjust the default thresholds to suit their target performance values.

**Supplementary information:** Supplementary data are available online.

## Introduction

SpliceAI is a neural network that predicts splicing from a pre-mRNA sequence (Jaganathan et al., 2019). Previous evaluations (Ha, Kim, & Jang, 2021; Moles-Fernández et al., 2021; Riepe, Khan, Roosing, Cremers, & ’t Hoen, 2021; Rowlands et al., 2021; Wai et al., 2020) have identified SpliceAI as the best predictor of variants that impact splicing, here termed spliceogenic variants. These studies assessed SNVs and small indels across multiple locations (i.e. splice site motifs, deep intronic regions >20 bp from the acceptor and >6 bp from the donor site, and exonic). They used the maximum delta score (of the four possible output scores) that passed the respective study-designated thresholds to predict variant spliceogenicity, but did not assess the splicing aberration type.

SpliceAI sensitivity for detecting spliceogenic intronic variants >50 bp from exons was originally reported to be 41% using a 0.5 maximum delta score threshold (Jaganathan et al., 2019), but an improved sensitivity of 94% was observed for variants >20 bp from exons by lowering the threshold to 0.05 (Moles-Fernández et al., 2021). Paired donor-acceptor splice site scores were observed for validated pseudoexonization events (Moles-Fernández et al., 2021). Moreover, manual checking of donor-acceptor splice site pairing was incorporated into a scheme to prioritize likely spliceogenic deep intronic variants (Qian et al., 2021).

We developed the SAI-10k-calc to systematically predict different SNV-associated splicing aberrations, altered transcript sizes, and consequent amino acid sequences, with a focus on accurate prediction of aberration sizes due to deep intronic variation.

## Methods

SAI-10k-calc was designed to predict specific types of splicing aberrations, namely: pseudoexonization, partial intron retention, partial exon deletion, (multi)exon skipping, and whole intron retention. Its features were derived from the application of all four raw delta scores and their corresponding delta positions generated by the SpliceAI tool (Jaganathan et al., 2019) using the maximum distance of +/− 4999 bp flanking the variant of interest. SAI-10k-calc can process SpliceAI scores resulting from SNVs at any exonic or intronic position, but not scores resulting from indels due to the complexity of distance interpretations for such variants. The decision flowchart is shown in Supplementary File 1.

We established default thresholds for SpliceAI delta scores (0.02–0.2 for exon skipping or whole intron retention and 0.02–0.05 for pseudoexon gain) and the gained exon size range of 25–500 bp based on two training sets derived from published splicing data: (1) SNVs in *BRCA1, BRCA2, MLH1, MSH2, MSH6*, and *PMS2* from Shamsani et al. (2019); and (2) deep intronic SNVs in various Mendelian disease genes from Moles-Fernández et al. (2021) (Supplementary Table S1). The 0.2 upper threshold for exon skipping is based on the lower limit set by SpliceAI developers (Jaganathan et al., 2019). For deep intronic variants, the 0.05 upper threshold for pseudoexon gain is also supported by previous findings (Moles-Fernández et al., 2021). The 25–500 bp exon size range encompasses the optimal size for efficient splicing that is between 50–250 bp (Movassat, Forouzmand, Reese, & Hertel, 2019) and is expected to capture most gained pseudoexons.

### Usage and features

SAI-10k-calc (R code version) requires two input files: a SpliceAI output VCF file and a tab-separated file with gene names and RefSeq transcript IDs (to match transcripts used in SpliceAI calculations). SAI-10k-calc was developed using human genome reference GRCh37, but is compatible with GRCh38.

The SAI-10k-calc output is a tab-separated file with summary of splicing predictions indicating the type of aberration, possible combinations of aberrations (e.g. one SNV resulting in both exon skipping and partial intron retention), the exact size of inserted and/or deleted sequences, and the effect on reading frame and translation (Figure 1). The latter is critical to predict the pathogenicity of the splicing alteration, and to design and interpret laboratory validation experiments. Amino acid sequence predictions could also be useful for additional applications, for example cancer neoantigen predictions (Yarchoan, Johnson, Lutz, Laheru, & Jaffee, 2017).

**Figure 1.**
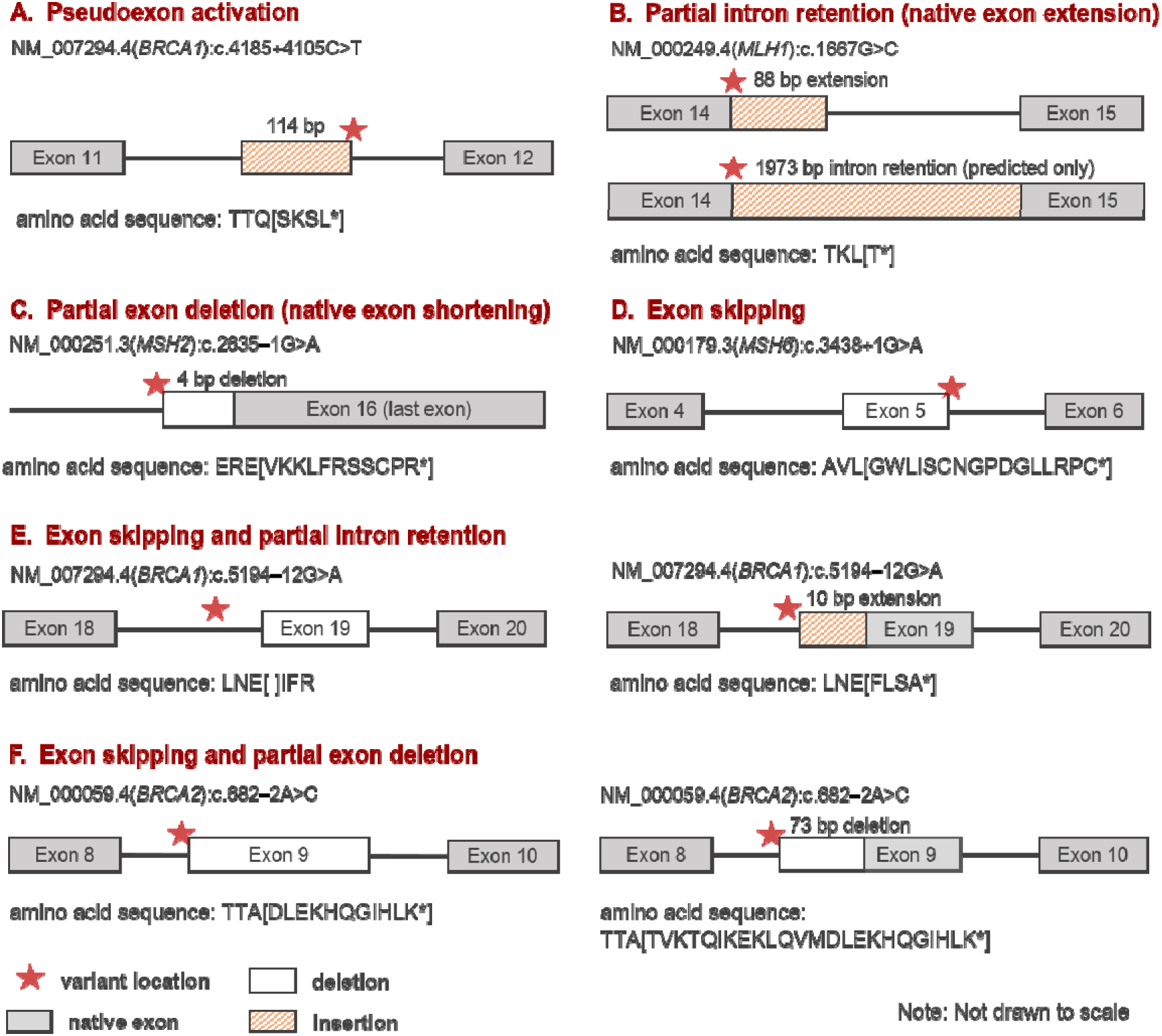
Types of splicing aberrations predicted by the SpliceAI-10k calculator. Six SNVs that were experimentally confirmed to alter splicing are correctly predicted by SAI-10k-calc (A to F). Of these, two represent correct prediction of combinations of splicing aberration types (E, F). Amino acid sequence predictions include three amino acids preceding the first variant amino acid, followed by the modified sequence inside square brackets. In-frame amino acid deletions (example in panel E) are indicated by blank square brackets flanked by three wild type amino acids preceding and following the deleted sequence.

We also provide a lightweight Microsoft Excel spreadsheet (Supplementary File 2, processing up to 1000 SNVs) that predicts the types and sizes of aberrations. In this version, users need to provide the raw scores generated by either SpliceAI Lookup (https://spliceailookup.broadinstitute.org/) or SpliceAI run from the command line. However, the predicted aberration sizes for partial intron retentions or partial exon deletions for this lightweight version are less accurate than the R code implemented version. Specifically, the R code uses native splice site positions derived from the given RefSeq transcript, whereas the lightweight version uses the SpliceAI-predicted acceptor and donor site positions. For example, the Excel version incorrectly predicted NM_007294.4(*BRCA1*):c.4868C>G to result in a 125-bp partial exon deletion, while the R code gave the correct size of 119 bp.

We note that, due to SpliceAI limitations, the calculator cannot be designed to predict three specific combinations of aberrant transcripts: (1) exon skipping and multi-exon skipping; (2) exon skipping and whole intron retention; and (3) partial exon deletion and partial intron retention. Multi-exon skipping and whole intron retention can only be predicted if the positions of donor and acceptor losses are within the analysis window, i.e., less than 4,999 bp from the variant.

### Performance

Using our training set data, SAI-10k-calc (R code version) has an overall sensitivity of 95% (441/464 confirmed spliceogenic SNVs) and specificity of 96% (715/748 non-spliceogenic SNVs) using our thresholds. Furthermore, SAI-10k-calc demonstrates high accuracy for prediction of pseudoexonization (85%), partial intron retention (84%), and exon skipping (81%), highlighting its applicability for prioritization of variants through clinical or research sequencing. R code output data from the training set variants are shown in Supplementary Table S2. General splicing prediction results and performance values are summarized in Supplementary Tables S3 and S4. Splicing aberration predictions (type and size) are detailed in Supplementary Tables S5-8.

## Acknowledgement

Emma Tudini and Vaishnavi Nathan of the QIMR Berghofer Molecular Cancer Epidemiology Laboratory updated the *BRCA1* and *BRCA2* splicing table that partly provided curated splicing data for training set 1.

