## Supplementary File 1 for "SpliceAI-10k calculator for the prediction of pseudoexonization, intron retention, and exon deletion"

**SpliceAI-10k calculator for the prediction of pseudoexonization, intron retention, and exon deletion**

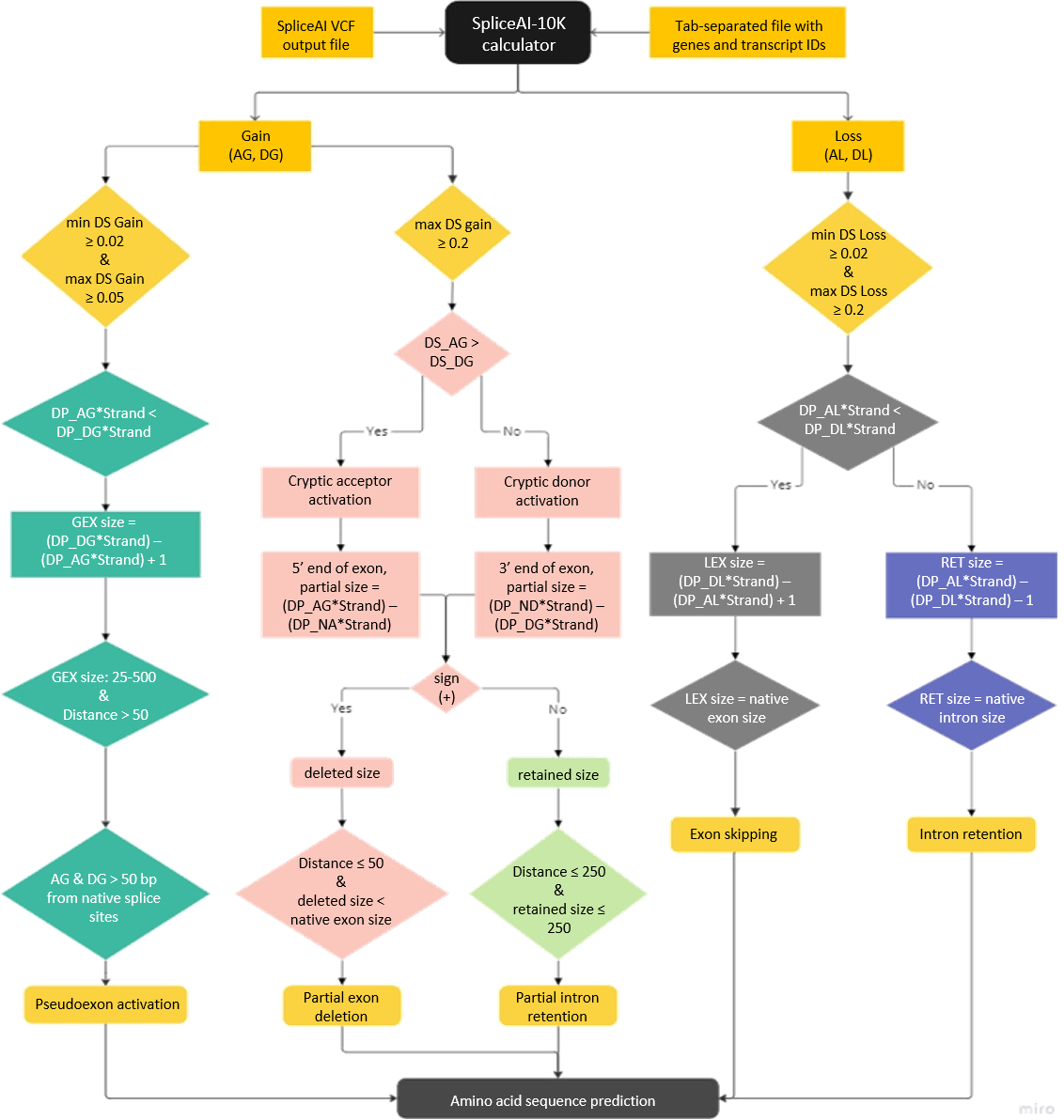

**SpliceAI-10k calculator decision flowchart.** An intronic location threshold of >250 bp was set as an exclusive region for pseudoexonization. An intronic region within 50 bp of the exon-intron junction was set as a threshold for SNVs with potential to cause exon skipping. An intronic location within 250 bp of the exon-intron junction was set as the threshold for SNVs that lead to multiple aberration types (i.e. combinations of pseudoexonization, partial intron retention, and exon skipping). Abbreviations are spelled out below.

| **List of Abbreviations** | |
| --- | --- |
| DS_AG | delta score for acceptor gain |
| DS_AL | delta score for acceptor loss |
| DS_DG | delta score for donor gain |
| DS_DL | delta score for donor loss |
| DP_AG | delta position for acceptor gain |
| DP_AL | delta position for acceptor loss |
| DP_DG | delta position for donor gain |
| DP_DL | delta position for donor loss |
| DP_NA | delta position of the native acceptor in the given transcript |
| DP_ND | delta position of the native donor in the given transcript |
| Distance | variant intronic distance (bp) from the nearest exon-intron junction |
| GEX | gained exon |
| LEX | lost exon |
| RET | retained intron |
| Strand | forward (1) or reverse (-1) strand orientation |
| VCF | variant call format |

**SPLICEAI-10K CALCULATOR OPTIONS**

**Gain, delta score threshold:**

DS_AGDG_MIN (default ≥ 0.02)

DS_AGDG_MAX (default ≥ 0.05)

For pseudoexonization, one splice site must have a score that passes the maximum delta score threshold (DS_AGDG_MAX) for splice site gain, and its splice site pair must have a score greater than or equal to the lowest acceptable delta score (DS_AGDG_MIN).

**Gained exon size range:**

GEX_size_MIN (default ≥ 25)

GEX_size_MAX (default ≤ 500)

Gained exon size must be within the range bounded by the lower size limit (GEX_size_MIN) and upper size limit (GEX_size_MAX).

**Loss, delta score threshold:**

DS_ALDL_MIN (default ≥ 0.02)

DS_ALDL_MAX (default ≥ 0.2)

For exon skipping or whole intron retention, one splice site must have a score that passes the maximum delta score threshold (DS_ALDL_MAX) for splice site loss, and its splice site pair must have a score greater than or equal to the lowest acceptable delta score (DS_ALDL_MIN).

**Cryptic splice site activation score threshold:**

DS_AG (default ≥ 0.2)

DS_DG (default ≥ 0.2)

Created/activated cryptic acceptor or donor splice site must pass their respective delta score threshold (DS_AG or DS_DG).

The 0.2 upper threshold for exon skipping, whole intron retention, and cryptic splice site activation is based on the lower limit set by the developers of SpliceAI (Jaganathan et al., 2019). For deep intronic variants, the 0.05 upper threshold for pseudoexon gain is supported by the findings of Moles-Fernández et al. (2021). The 25–500 bp exon size range encompasses the optimal size for efficient splicing that is between 50–250 bp (Movassat, Forouzmand, Reese, & Hertel, 2019) and expected to capture most gained pseudoexons.

More information can be found at <https://github.com/adavi4/SAI-10k-calc>.
